# Kinetic Signatures of RAS–BRAF Engagement Reveal Isoform Selectivity, Oncogenic Amplification, and Mutation-Dependent Inhibitor Resistance

**DOI:** 10.64898/2026.05.24.727535

**Authors:** Sravani Malasani, Zhihong Wang

## Abstract

The RAS-RAF-MEK-ERK (MAPK) signaling cascade is a central regulator of cellular proliferation and differentiation, and its dysregulation is a frequent driver of oncogenesis. RAF activation requires recruitment by GTP-bound RAS at the plasma membrane, however, the precise molecular determinants governing RAS-RAF engagement remain incompletely understood. Here, we show that the BRAF N-terminal region forms a cooperative BSR-CRD autoinhibitory gate that restricts RAS engagement and encodes isoform-specific kinetic behavior. Using OpenSPR and BLI, together with NanoBiT cellular assays, we reveal a kinetic encoding mechanism in which KRAS and NRAS define distinct regimes of BRAF engagement: KRAS exhibits stability-driven, long-lived complex formation, whereas NRAS displays frequency-driven, transient interactions. Oncogenic mutations reshape these regimes by selectively stabilizing RAS-BRAF association without uniformly increasing affinity, amplifying isoform-specific kinetic signatures. Pharmacological profiling further reveals isoform-dependent sensitivity of RAS-RAF disruption governed by nucleotide state and compatibility with the BSR-CRD gate. Together, these findings establish that KRAS and NRAS operate through distinct kinetic regimes of BRAF engagement governed by a structurally gated N-terminal regulatory architecture encoding temporal and pharmacological specificity in MAPK signaling.

**Significance:** This study defines a structural and kinetic framework governing RAS-BRAF engagement in MAPK signaling. We define the BRAF BSR-CRD region as a cooperative autoinhibitory gate that controls RAS accessibility and encodes isoform-specific interaction dynamics. KRAS and NRAS exhibit distinct kinetic regimes of BRAF engagement, which are further reshaped by oncogenic mutations. We further show that inhibitor sensitivity is strongly influenced by nucleotide state and isoform-specific Switch II pocket architecture. Together, these findings establish kinetic encoding as a key determinant of RAF activation and therapeutic response in RAS-driven cancers.

## Introduction

RAS and RAF proteins are key components of the mitogen-activated protein kinase (MAPK) signaling pathway, which regulates cell growth, division, differentiation, and survival^1^. RAS, a small GTPase, acts as a molecular switch, cycling between an inactive GDP-bound state and an active GTP-bound state^2^, a process regulated by guanine nucleotide exchange factors (GEFs) and GTPase-activating proteins (GAPs)^3^. RAF, a serine/threonine protein kinase, is autoinhibited in the cytosol and becomes activated once recruited to the membrane by activated RAS, where it dimerizes and activates downstream signaling through MEK and ERK^4–6^. Activated ERK in turn phosphorylates a diverse set of nuclear transcription factors to drive cellular responses.

The RAF kinase family (ARAF, BRAF, and CRAF) shares a conserved domain organization consisting of CR1, CR2, and CR3^7^. In the inactive state, interactions between the N-terminal regulatory region (CR1-CR2) and the C-terminal kinase domain (CR3) maintain RAF in an autoinhibited conformation^7^. Activation occurs following RAS-mediated conformational rearrangements and membrane recruitment that promote RAF dimerization and kinase activation. CR1 contains the RAS-binding domain (RBD) and the cysteine-rich domain (CRD), which cooperatively facilitate interaction with activated RAS and mediate the recruitment of RAF to the plasma membrane^2,7^. While the RBD drives high-affinity nanomolar binding to RAS, the CRD contributes additional contacts that enhance RAF-RAS association and is required for efficient signaling despite its weaker intrinsic affinity^8–11^. Structural studies of CRAF-RAS complexes have defined CRD contacts with KRAS/HRAS that are essential for activation, including interactions with the interswitch region and helix α5 of RAS^12^. Cryo-EM structures of autoinhibited BRAF with 14-3-3 and MEK showed that CRD-kinase domain contacts stabilize inactive monomers, with 14-3-3 blocking dimerization^7, 13^.The CR2 region contains regulatory phosphorylation sites, including BRAF S365 and CRAF S259, which serve as binding sites for 14-3-3 proteins and contribute to maintaining RAF in an autoinhibited state^7^. The CR3 region comprises the kinase domain responsible for the phosphorylation of downstream substrate MEK upon release of autoinhibitory interactions and subsequent dimerization, thereby propagating MAPK signaling^6, 14^.

Among the three RAF isoforms, BRAF exhibits the highest basal kinase activity and is the most frequently mutated RAF family member in human cancers^15, 16^. The most prevalent BRAF alterations are mutations such as V600E/K, which led to the development of first-generation BRAF inhibitors and have transformed the clinical landscape for melanoma. Moreover, BRAF displays distinct regulatory behavior compared with CRAF and ARAF, including stronger CRD-mediated autoinhibition and isoform-specific RAS preferences^15–18^. Central to these isoform-specific constraints is the N-terminal BRAF-specific region (BSR), a structural motif unique to BRAF. Recent biochemical characterization reveals that the BSR acts in tandem with the adjacent cysteine-rich domain (CRD) to exert a negative regulatory constraint, maintaining autoinhibition^19^ and attenuating unauthorized CRD-lipid interactions with the plasma membrane^20^. Furthermore, the BSR is believed to form electrostatic attractions with the polybasic hypervariable region (HVR) of KRAS rather than HRAS^12^.

The three primary RAS genes encode four RAS protein isoforms, including HRAS, NRAS, with KRAS generating two splicing variants, KRAS4A and KRAS4B, through alternative RNA splicing. RAS proteins, which exhibit approximately 90% sequence identity across isoforms, consist of a highly conserved globular (G) domain (residues 1-166) and a C-terminal hypervariable region (HVR, residues 166-188/189)^3, 21^. The G domain contains two dynamic regions, Switch I (residues ∼30-38) and Switch II (residues ∼60-76), which undergo conformational rearrangements upon GDP-GTP exchange and directly regulate effector binding as well as GTP hydrolysis^22–24^. These switch regions form the principal interface for effector engagement, including RAF binding, and are essential for propagating downstream signaling. HVR undergoes isoform-specific post-translational modifications, including farnesylation and palmitoylation, and contains polybasic motifs that collectively govern membrane association and subcellular localization^25, 26^.

Among RAS family members, KRAS is the most frequently mutated isoform, accounting for 85% of all RAS-mutant tumors, compared with 11% for NRAS and 4% for HRAS^27^. These mutations primarily occur at three hotspot codons: G12, G13, and Q61^28^. Notably, RAS isoforms display distinct mutation frequencies across the three hotspots, with G12 mutations being the most prevalent in KRAS, whereas Q61 represents the predominant mutational hotspot in both NRAS and HRAS^29^. As the least studied RAS isoform, the contribution of NRAS to MAPK hyperactivation is frequently overlooked, yet oncogenic NRAS alterations are a primary driver of acquired resistance to first-generation BRAF inhibitors^30^, making a deeper biochemical understanding of the NRAS-RAF interface essential for designing next-generation combination strategies. The discovery of covalent inhibitors Sotorasib (AMG510)^31^ and Adagrasib (MRTX849)^32^ targeting the KRAS G12C mutant, which selectively bind the switch II pocket and trap KRAS in its inactive GDP-bound state, has shifted the paradigm of RAS being regarded as “undruggable”. More recently, next-generation KRAS inhibitors with distinct mechanism of action have expanded the therapeutic landscape, including inhibitors targeting non-G12C KRAS mutants, such as BI2865^33^, a highly selective inhibitor of pan KRAS(GDP-bound), and RMC-6236^34^, a pan-RAS(ON) inhibitor that targets active RAS proteins across multiple mutant isoforms^35^.

Despite extensive study of RAS-RAF signaling, several key mechanistic questions remain unresolved. Specifically, it is unclear how oncogenic RAS mutations alter the conformational landscape of the BRAF N-terminal regulatory region beyond the canonical RBD interface. NRAS remains a major oncogenic driver, particularly in melanoma and hematologic malignancies, yet remains comparatively less explored in therapeutic targeting, underscoring the need for deeper mechanistic understanding of NRAS-specific signaling regulation. Furthermore, while multiple classes of RAS-targeted inhibitors have been developed, their direct impact on RAS-RAF complex formation, including binding kinetics, isoform dependence, mutation-specific effects, and consequences for RAF engagement, is insufficiently characterized. To address these limitations, we combined biophysical and cellular approaches to systematically interrogate how distinct oncogenic states and pharmacological interventions modulate RAS-RAF assembly across various isoforms.

## Results

### NRAS binds the BRAF N-terminal domain with intermediate affinity relative to KRAS and HRAS

Although the binding affinities of BRAF N-terminal domain variants with HRAS and KRAS have been previously elucidated by our group^19^, a critical next step was to determine how NRAS interacts with the BRAF N-terminus. To investigate this, we purified four N-terminal BRAF constructs, each tagged with MBP at the N-terminus: NT1 (BSR+RBD+CRD; amino acids 1-288), NT2 (BSR+RBD; 1-227), NT3 (RBD+CRD; 151-288), and NT4 (RBD; 151-227). GMPPNP- loaded NRAS and KRAS, as well as their inactive counterparts, were also purified (Figure 1A & B). We used OpenSPR to characterize these binding interactions between NRAS and these diverse BRAF regulatory domains.

**Figure 1:**
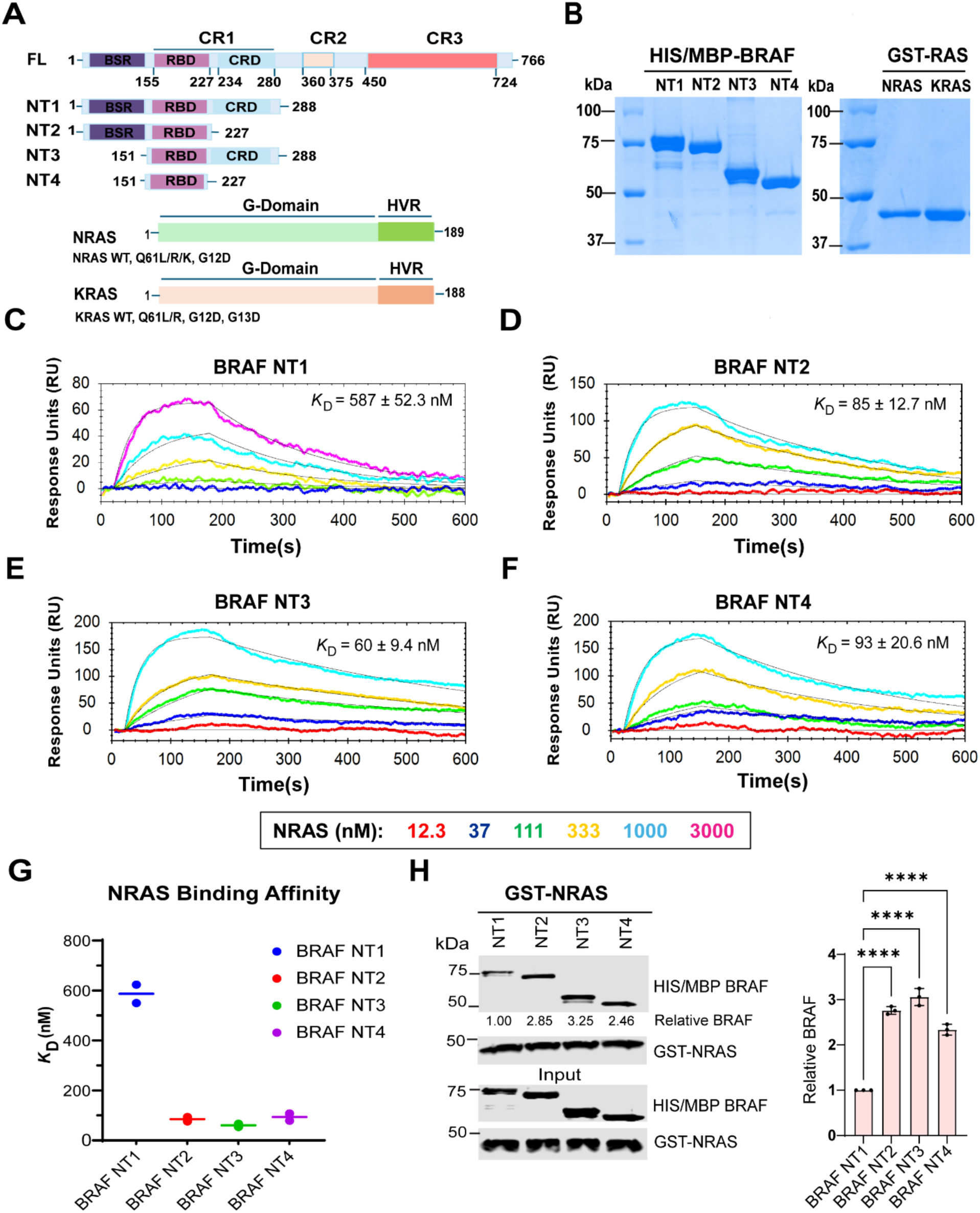
Domain architecture of the BRAF N-terminal region differentially regulates NRAS binding. A) Schematic representation of BRAF N-terminal constructs used in this study, including NT1 (BSR-RBD-CRD), NT2 (BSR-RBD), NT3 (RBD-CRD), and NT4 (RBD). Not shown:6Xhis/MBP tag on N-terminal of NT proteins and GST tag on N-terminal of KRAS. BSR: BRAF-specific region; RBD: RAS binding domain; CRD: cysteine rich domain. B) Coomassie-stained SDS-PAGE gels showing purified BRAF NT1-4 along with KRAS and NRAS. C-F) OpenSPR sensorgrams of NRAS binding to BRAF NT1-4 immobilized on NTA sensors. Experimental data are shown with the corresponding global fits generated using a 1:1 fitting model. G) Summary diagram showing the binding affinity of NRAS for BRAF N-terminal constructs (NT1–NT4). Dissociation constants (*K*_D_, nM) were determined by independent binding assays. Each point represents replicate measurements. H) Western blot of GMPPNP-bound NRAS pulled down on glutathione resin to probe for BRAF NT1-4 binding. Band intensities were quantified by densitometry and normalized to control conditions. Bar graphs were generated and statistical analysis was performed using GraphPad Prism using ordinary one-way ANOVA. Data are presented as mean ± SD from three independent experiments (n = 3).

The full N-terminal construct, BRAF NT1, exhibited relatively weak binding to NRAS, with a dissociation constant (*K*_ᴅ_) of 587 ± 52.32 nM (Figure 1C). Kinetic analysis revealed a slow association rate (*k*_on_ = 8.08×10³ M^−1^ s^−1^) and a faster dissociation rate (*k*_off_ = 4.66×10^-3^ s^−1^), indicating inefficient complex formation and reduced stability of the NRAS-BRAF NT1 complex. In comparison, this interaction was substantially weaker than the previously reported binding of KRAS to NT1 (*K*_ᴅ_ = 267 ± 9.89 nM)^19^.

To evaluate the contribution of individual regulatory domains, we next examined BRAF NT2, which lacks the CRD. Removal of the CRD resulted in a marked, nearly seven-fold increase in binding affinity, with NT2 bound NRAS with a *K*_ᴅ_ of 85 ± 12.7 nM (Figure 1D). This improvement was driven by a substantially faster association rate (*k*_on_ = 3.24×10^4^ M^−1^ s^−1^) coupled with a reduced dissociation rate (*k*_off_ = 2.79×10^-3^ s^−1^). These kinetic changes indicate more efficient complex formation and improved stability upon removal of the CRD, suggesting that the full N-terminal architecture may partially restrain RAS engagement. Despite this increased affinity, NT2-NRAS binding remained weaker than interactions reported for NT2-HRAS (7.5 ± 3.5 nM) and NT2-KRAS (33 ± 5 nM)^19^.

We further analyzed BRAF NT3 (RBD-CRD) to determine the contribution of the BSR domain. NT3 bound NRAS with a *K*_ᴅ_ of 60.3 ± 9.47 nM (Figure 1E), representing a modest improvement over NT2. Kinetic analysis revealed a further increase in the association rate (*k*_on_ = 4.17 ×10^4^ M^−1^ s^−1^) while maintaining a similarly slow dissociation rate (*k*_off_ = 2.54×10^-3^ s^−1^). Under these conditions, NRAS displayed an affinity comparable to KRAS (86.5 ± 35 nM) but weaker than HRAS (22 ± 11 nM)^19^.

Finally, to isolate the specific contribution of CRD, we tested BRAF NT4, which contains only the RBD domain. NT4 bound NRAS with a *K*_ᴅ_ of 93.45 ± 20.57 nM (Figure 1F). The kinetic parameters revealed a rapid association rate (*k*_on_ = 4.28 ×10^4^ M^−1^ s^−1^) but a slightly faster dissociation rate (*k*_off_ = 3.78 ×10^-3^ s^−1^) compared with NT3. This suggests that while the RBD is sufficient to mediate strong binding, the presence of adjacent CRD modestly enhances complex stability. In comparison, HRAS exhibited stronger binding (19 ± 11 nM), whereas KRAS showed intermediate affinity (52 ± 22.62 nM)^19^.

A summary of the *K*_ᴅ_ values for NRAS interactions with BRAF N-terminal constructs (NT1-NT4) is shown in Figure 1G; Kinetic parameters are summarized in Table S1, highlighting the substantially weaker interaction observed for NT1 and the robust binding exhibited by NT2, NT3, and NT4. These results indicate that the RBD serves as the primary determinant of RAS binding, whereas both BSR-RBD and RBD-CRD exhibited enhanced binding affinity relative to RBD alone. However, the BSR-RBD-CRD construct showed significantly reduced binding affinity, indicating that simultaneous incorporation of BSR and CRD negatively affected NRAS engagement. Consistent with these biophysical measurements, pull-down assays revealed weak binding between NT1 and NRAS, while NT2, NT3, and NT4 displayed stronger interactions (Figure 1H), confirming the critical role of the RBD containing constructs in mediating stable RAS-BRAF engagement.

### Codon 61 mutations in NRAS strengthen interaction with the BRAF N-terminus

To determine how oncogenic NRAS mutations influence its interaction with BRAF, we measured the binding affinities of several cancer-associated mutants (Q61L, Q61R, Q61K, and G12D) toward the BRAF NT1 construct using OpenSPR. Wild-type NRAS displayed relatively weak binding to BRAF NT1, with a dissociation constant (*K*_ᴅ_) of 587 ± 52.32 nM. In contrast, all four mutant forms of NRAS demonstrated significantly enhanced binding affinities (Figure 2A-D). Notably, the Q61L and Q61R mutants showed the most robust interaction, with *K*_ᴅ_ values of 87 ± 6 nM and 91 ± 8 nM, reflecting a ∼7-fold increase in affinity compared with wild-type NRAS. Similarly, the Q61K and G12D mutants exhibited *K*_ᴅ_ values of 116 ± 28 nM and 116 ± 2.5 nM, corresponding to roughly a 5-fold improvement in binding affinity. Kinetic analysis of those NRAS mutants demonstrated a consistent increase in complex formation efficiency, as indicated by uniformly elevated association rates (*k*_on_ = 3.35 ×10^4^ - 4.30 ×10^4^ M^−1^ s^−1^). In contrast, dissociation rates remained relatively similar across all variants (*k*_off_ = 3.26 × 10^−3^ - 3.96 × 10^−3^ s^−1^). A summary of the *K*_ᴅ_ values for all NRAS variants interacting with BRAF NT1 is presented in Figure 2E while the complete kinetic parameters are listed in Table S2. Together, these data demonstrate that oncogenic NRAS mutants substantially enhance binding affinity toward the BRAF N-terminus relative to wild-type NRAS.

**Figure 2.**
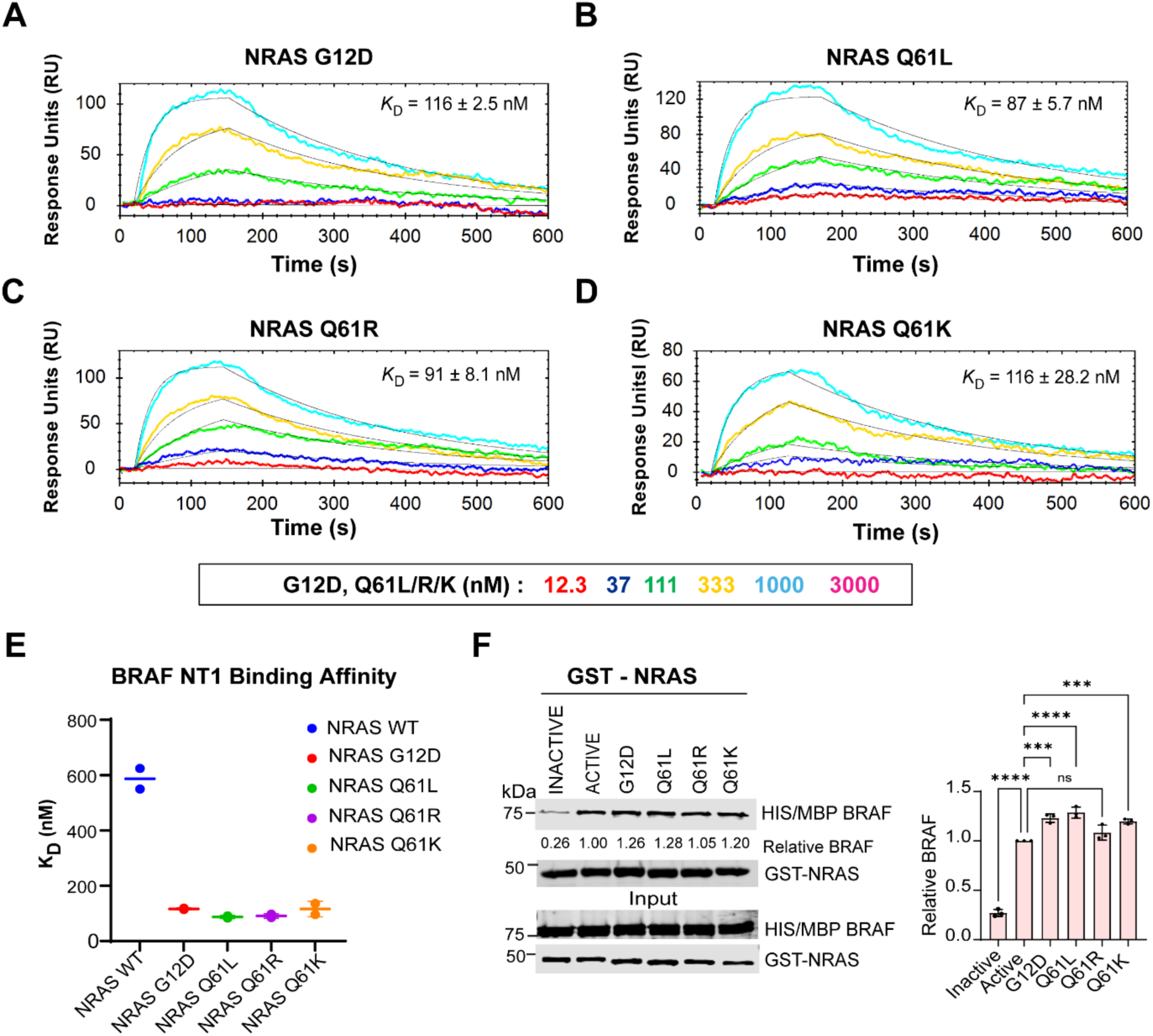
Oncogenic NRAS mutations enhance binding to the BRAF N-terminus. A-D) OpenSPR sensorgrams showing binding interactions between BRAF NT1 (BSR-RBD-CRD) and NRAS variants, including wild-type and mutants (Q61L, Q61R, Q61K, and G12D). E) Summary of dissociation constants (*K*_D_, nM) for NRAS variants interacting with BRAF NT1. Each point represents the mean ± SD from replicate measurements. F) GST pull-down assays comparing interactions of BRAF NT1 with GDP-bound NRAS, GMPPNP-loaded NRAS, and NRAS mutants. Band intensities were quantified by densitometry, normalized to control, and analyzed using GraphPad Prism with ordinary one-way ANOVA. Data are shown as mean ± SD from three independent experiments (n = 3). All OpenSPR experimental data were analyzed with the corresponding global fits generated using a 1:1 fitting model.

To further validate these biophysical observations, we performed GST pull-down assays using NRAS in both its inactive GDP-bound and active GMPPNP-bound states, along with the GMPPNP-bound NRAS mutants (Figure 2F). Consistent with the OpenSPR measurements, GDP-loaded NRAS showed minimal interaction with BRAF NT1, whereas GMPPNP-loaded NRAS and all four oncogenic mutants exhibited markedly stronger binding. These results confirm that the active form of NRAS, particularly oncogenic mutant variants, engages the BRAF N-terminus more robustly.

### Oncogenic KRAS mutations significantly enhance association with the BRAF N-terminus

Following characterization of NRAS interactions with the BRAF N-terminus, we next examined whether oncogenic mutations in KRAS similarly influence binding to BRAF NT1. Binding kinetics of several clinically relevant KRAS variants (G12D, G13D, Q61R, and Q61L) were measured using OpenSPR (Figure 3A-D). Wild-type KRAS bound BRAF NT1 with a dissociation constant (*K*_ᴅ_) of 267 ± 9.8 nM (Figure 3E; Table S3). Consistent with the patterns observed for NRAS, substitutions at codon 61 and codon 13 significantly enhanced the KRAS-BRAF NT1 interaction. KRAS (G13D) and KRAS (Q61L) exhibited *K*_ᴅ_ values of 24.8 ± 1.97 nM and 19.7 ± 1.34 nM, respectively, corresponding to approximately 11-fold and 13-fold increases in binding affinity relative to wild-type KRAS. Similarly, the KRAS(Q61R) mutant displayed enhanced binding, with a *K*_ᴅ_ of 37.7 ± 15.7 nM, representing an approximately 7-fold increase in affinity. In contrast, the G12D substitution produced only a modest effect, with a *K*_ᴅ_ of 203.5 ± 14.8 nM. Biochemical pull-down assays supported these kinetic observations. A broader panel of oncogenic KRAS variants, including G13D, Q61H, Q61R, and Q61L, showed enhanced association with BRAF NT1 compared with wild-type KRAS, whereas G12D and G12V variants displayed similar association compared with wild-type KRAS (Figure 3F). Collectively, these findings demonstrate that oncogenic mutations in RAS, particularly those at codon 61 and codon 13, substantially strengthen interactions with the BRAF N-terminal region.

**Figure 3:**
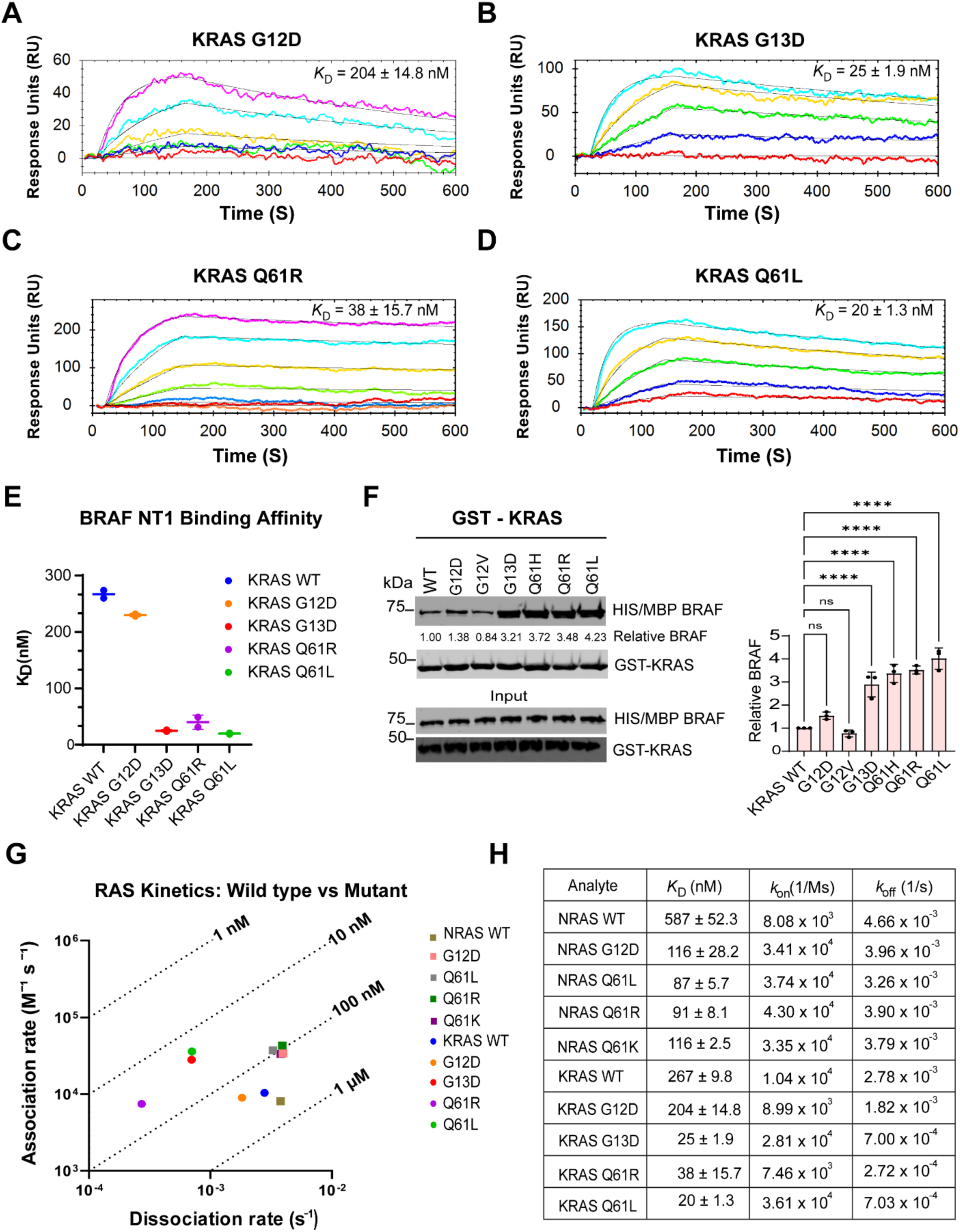
Oncogenic KRAS mutations enhance binding affinity and kinetic stability of the KRAS-BRAF interaction. A-D) OpenSPR sensorgrams showing binding kinetics of BRAF NT1 (BSR–RBD–CRD) with wild-type (WT) KRAS and oncogenic KRAS variants (G12D, G13D, Q61R, and Q61L). E) Summary of dissociation constants (*K*_D_, nM) for WT and mutant KRAS proteins binding to BRAF NT1. Each point represents the mean ± SD from replicate measurements. F) GST pull-down assays comparing interactions of a broader panel of oncogenic KRAS variants (G12D, G12V, G13D, Q61H, Q61R, and Q61L) with WT KRAS towards BRAF NT1. Band intensities were quantified by densitometry, normalized to control, and analyzed by one-way ANOVA in GraphPad Prism (n = 3, mean ± SD). G) Kinetic scatter plot comparing association (*k_on_*) and dissociation (*k_off_*) parameters for RAS variants interacting with BRAF NT1. H) Summary of kinetic parameters (*k_on_, k_off_, and K*_D_) obtained from OpenSPR kinetic fitting with BRAF NT1 as the ligand and RAS mutants as the analytes. All OpenSPR experimental data are analyzed with the corresponding global fits generated using a 1:1 fitting model.

To further elucidate the biophysical basis underlying these interactions, we analyzed the individual kinetic parameters using a kinetic scatter plot (Figure 3G). The improved affinities of KRAS variants were driven by a combination of faster association rates (*k*_on_) and significantly stabilized dissociation kinetics (*k*_off_). Notably, comparative analysis between the two RAS isoforms revealed a distinct kinetic signature for KRAS mutants. While oncogenic NRAS variants (Q61L, Q61R, Q61K, G12D) displayed relatively rapid dissociation from BRAF NT1 (∼ 3.3 - 4.0×10^−3^ s^−1^), the corresponding KRAS mutants (Q61R, Q61L, G13D) exhibited markedly slower dissociation rates (∼ 2.7 - 7.0×10^-4^ s^−1^). These findings suggest that oncogenic KRAS mutations confer greater kinetic stability to the RAS-BRAF complex than the corresponding NRAS mutations.

### RAS mutations reshape BRAF engagement and inhibitor sensitivity in cells

We next assessed RAS-BRAF interactions in HEK293 cells using a NanoBiT luminescence complementation assay, a well-established method for quantifying protein interactions in live cells. In this system, RAS is fused to the larger BiT subunit, while BRAF is fused to the smaller BiT subunit. When RAS and BRAF interact, the resulting luminescence signal allows for the quantification of their binding intensity. To rule out the possibility that altered luminescence signals stem from changes in expression levels rather than differences in binding affinity, we quantified protein expression and found it remained constant across all compared conditions (Sup Fig 1A&B). Consistent with the biophysical data obtained using purified proteins, NanoBiT complementation assays revealed significantly increased luminescence for NRAS(Q61L/K)-BRAF and KRAS(G13D,Q61R)-BRAF pairs compared with their respective wild-type counterparts, indicating enhanced interaction in cells (Figure 4A & D).

**Figure 4:**
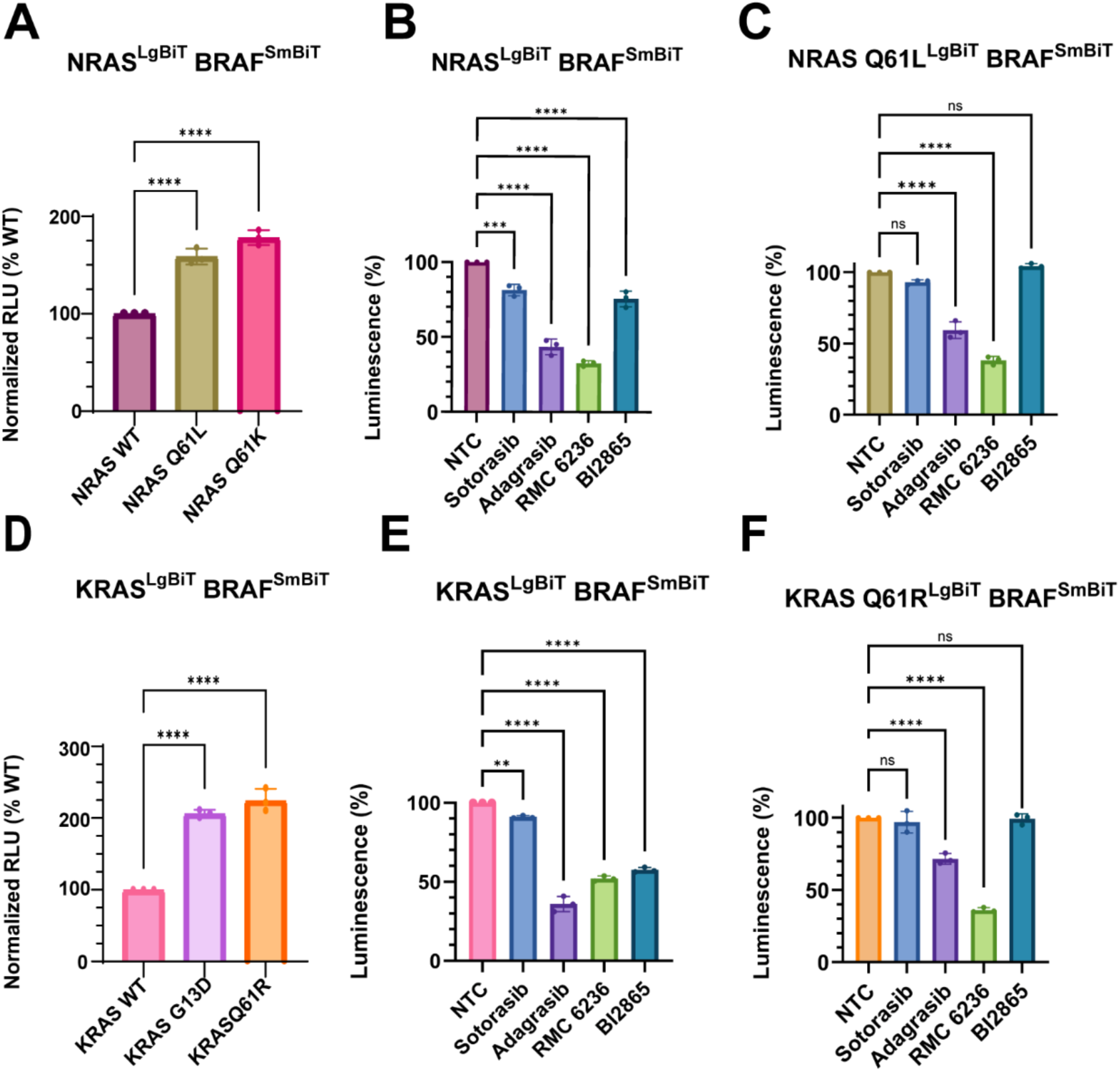
Quantification of RAS-BRAF interactions in response to oncogenic RAS mutations and RAS inhibitors in HEK293 cells. (A) NanoBiT complementation assay measuring NRAS-BRAF interaction in HEK293 cells. (B&C) Effects of RAS inhibitors (sotorasib, adagrasib, RMC6236, and BI2865) on WT NRAS-BRAF interactions (B) and NRAS Q61L-BRAF (C) interactions measured by NanoBiT assay. (D) NanoBiT complementation assay measuring KRAS-BRAF interaction in cells. (E&F) Effects of RAS inhibitors (sotorasib, adagrasib, RMC6236, and BI2865) on WT KRAS-BRAF interactions (E) and KRAS Q61R-BRAF (F) interactions measured by NanoBiT assay. All NanoBiT assay results were normalized to the untreated control condition and represent data collected from six independent biological replicates. Statistical analysis was performed using ordinary one-way ANOVA followed by Tukey’s honest significant difference (HSD) post-hoc test to evaluate multiple comparisons. Significance levels are indicated as *P < 0.05, **P < 0.01, and ***P< 0.001. Data are presented as mean ± standard deviation (SD), with individual points representing each biological replicate and corresponding *P*-values shown on the graphs.

To determine how pharmacological inhibition of RAS affects these interactions, HEK293 cells transiently transfected with NRAS-BRAF pairs were treated with several RAS inhibitors (Figure 4B). We quantified protein expression and confirmed that levels remained constant across all compared conditions (Sup Fig 2A&B). In cells expressing wild-type NRAS and BRAF, sotorasib and adagrasib reduced luminescence by approximately 18% and 56%, respectively, whereas RMC6236 produced the most pronounced reduction (∼67% inhibition). BI2865 resulted in a more modest decrease (∼24% inhibition). In cells expressing the NRAS(Q61L)-BRAF pair, sotorasib and adagrasib reduced luminescence by 7% and 40%, respectively, while RMC6236 again produced substantial inhibition (∼62% reduction). In contrast, BI2865 did not significantly alter luminescence under these conditions (Figure 4C). A similar trend was observed for the NRAS(Q61K)-BRAF interaction, where sotorasib and adagrasib resulted in 5% and 56% inhibition, respectively, while RMC6236 produced a strong reduction (∼75% inhibition). BI2865 again failed to produce a statistically significant effect (Sup Fig 3A).

A similar trend was observed for KRAS. In cells expressing WT KRAS-BRAF, treatment with adagrasib and RMC6236 resulted in the most pronounced displacement, with 64% and 48% inhibition of luminescence, respectively, Sotorasib showed minimal effect (9% inhibition), whereas BI-2865 induced a moderate reduction (42% inhibition) (Figure 4E). Protein expression levels were quantified and remained consistent across all conditions (Sup Fig 2C&D).

Consistent with our observations in the NRAS mutant context, KRAS(Q61R/G13D) selectively preserved effector engagement under inhibitor treatment. The Q61R mutation conferred complete resistance to sotorasib and BI2865 and significantly attenuated the efficacy of adagrasib (28% inhibition compared with 64% in WT KRAS) (Figure 4F; Sup Fig 3B). In contrast, the tricomplex RAS inhibitor RMC6236 maintained high potency against the Q61R variant, achieving 64% inhibition. Together, these data demonstrate that mutations at the G13 and Q61 hotspots consistently promote stronger BRAF NT1 recruitment through increased biochemical affinity and kinetic stability, while simultaneously creating allele-specific barriers to conventional switch-II pocket inhibitors.

### BLI analysis of KRAS and NRAS-BRAF NT1 interactions in the presence of Adagrasib and BI-2865

To further investigate how KRAS-targeted inhibitors differently modulate RAS-BRAF interactions, we quantified binding kinetics between KRAS or NRAS and BRAF NT1 using biolayer interferometry (BLI). NRAS and KRAS exhibited *K*_ᴅ_ values of 659 ± 68.59 nM and 210 ± 33.94 nM, respectively (Figure 5A &D). These values closely matched those obtained by OpenSPR (NRAS: 587 ± 52.32 nM; KRAS: 267 ± 9.89 nM), demonstrating strong agreement between the two biophysical measurements. The similar kinetic profiles obtained from BLI and OpenSPR further validate the utility of both methods for characterizing RAS–RAF protein interactions in solution.

**Figure 5:**
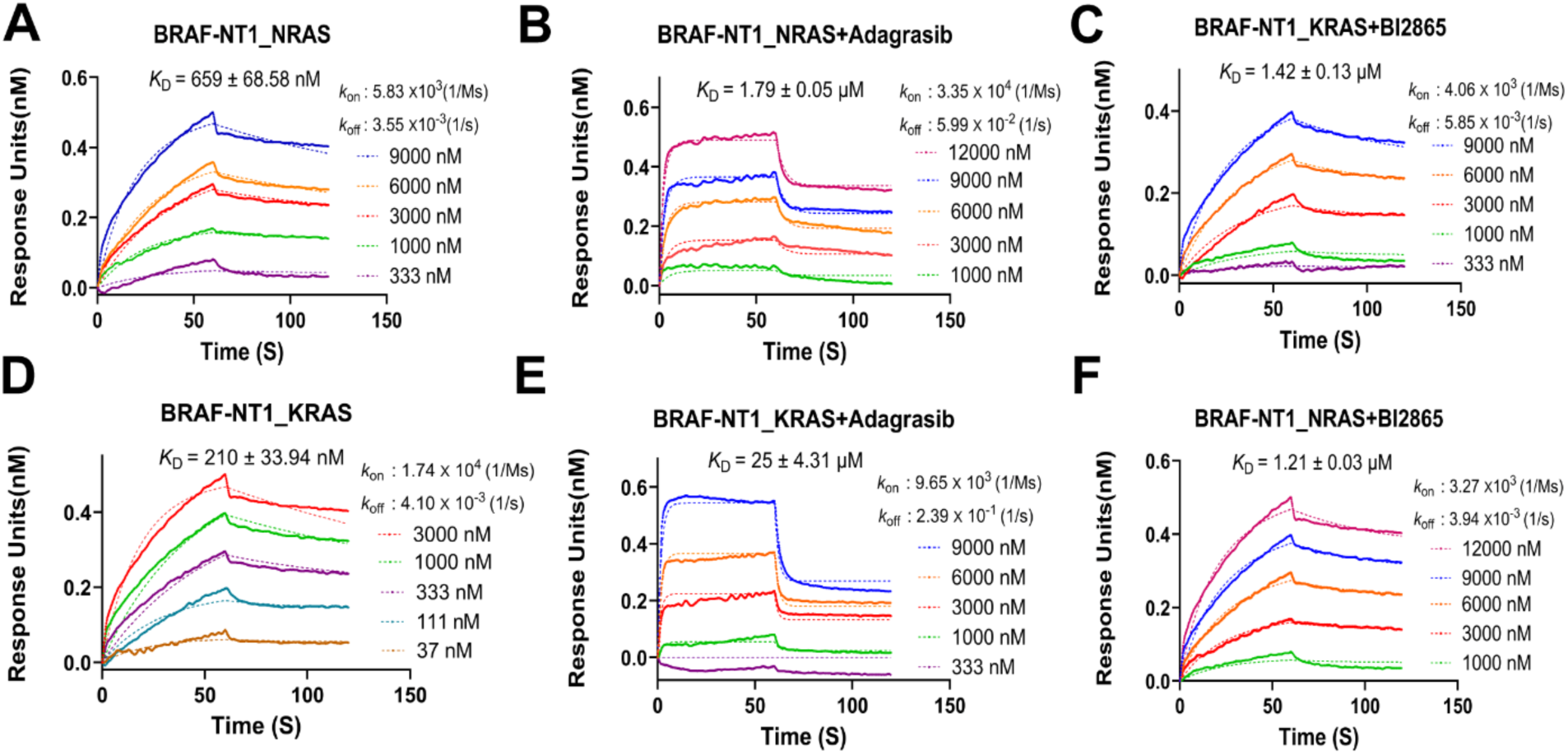
KRAS inhibitors distinctively disrupt the RAS–BRAF interactions. Biolayer interferometry (BLI) analysis of interactions between BRAF NT1 (BSR, RBD, CRD) and RAS. (A) Wild-type NRAS binding to BRAF NT1. (B-C) Wild-type NRAS binding in the presence of adagrasib or BI-2865, respectively. (D) Wild-type KRAS binding to BRAF NT1. (E-F) Wild-type KRAS-BRAF NT1 binding in the presence of adagrasib or BI-2865. RAS proteins were pre-incubated with 10 µM compounds for 2 h prior to BLI measurements. All BLI experiments were performed in triplicates. Sensorgrams were globally fitted to a 1:1 binding model to obtain the association (*k*_on_) and dissociation (*k*_off_) rate constants, and the binding affinity (*K*_D_) was calculated from the *k*_off_/*k*_on_ ratio. The fits showed good agreement with the data (R² > 0.9; χ² < 4).

Adagrasib treatment differentially affected the two RAS isoforms. KRAS-BRAF NT1 binding was strongly disrupted, with affinity decreasing from nanomolar to micromolar range (*K*_ᴅ_ = 25 ± 4.31 µM), corresponding to an approximately 120-fold reduction in binding strength (Figure 5E). This was accompanied by a pronounced increase in dissociation rate (*k*_off_ ∼ 2.39 × 10^−1^ s^−1^). In contrast, NRAS showed a more moderate reduction in affinity *K*_ᴅ_= 1.79 ± 0.05 µM) (Figure 5B), representing a ∼2.7-fold decrease, along with a smaller increase in dissociation rate (*k*_off_ ∼ 5.98 × 10^−2^ s^−1^), consistent with partial destabilization of the complex.

BI-2865 produced a more uniform attenuation of RAS-BRAF NT1 interactions across both isoforms (Figure 5C&F). KRAS binding was reduced to micromolar affinity (*K*_ᴅ_∼ 1.42 ± 0.13 µM; ∼ 6.8-fold decrease), while NRAS binding showed ∼1.8-fold decrease (*K*_ᴅ_ ∼ 1.21± 0.03 µM), reflecting modest reductions in binding affinity and complex stability across both isoforms. Across all datasets, kinetic fits were robust (R² > 0.97; Table S4 for full values), supporting the reliability of the derived kinetic parameters. Collectively, these results demonstrate isoform-dependent sensitivity to adagrasib and a modest disruption of RAS-BRAF NT1 engagement by BI-2865.

## Discussion

In this study, we systematically characterized interactions between the BRAF N-terminal regulatory domain and both wild-type NRAS and oncogenic N/KRAS isoforms using complementary biophysical and cellular approaches. By integrating OpenSPR, BLI, GST pulldown, and live-cell NanoBiT complementation assays, we identified isoform-and mutation-dependent differences in BRAF engagement, interaction kinetics, and inhibitor sensitivity. Collectively, our findings demonstrate that oncogenic RAS mutations, particularly those affecting codon 61, substantially enhance BRAF association while simultaneously altering susceptibility to switch-II pocket inhibitors.

Our data support a model in which the BRAF N-terminal regulatory assembly imposes conformational constraints on productive RAS engagement. Comparative analysis of the BRAF N-terminal constructs suggests that the BSR and CRD domains cooperatively modulate accessibility of the RBD-containing interaction surface. Although fusion of either BSR or CRD individually enhanced RAS binding relative to the isolated RBD, incorporation of both domains within the intact BSR-RBD-CRD assembly reduced overall affinity, indicating that the complete N-terminal regulatory module adopts a more constrained configuration. These observations are consistent with a structurally regulated mechanism in which productive RAS recruitment depends not only on the intrinsic affinity of the RBD but also on conformational accessibility within the larger N-terminal assembly.

Because BSR is unique to BRAF, this layer of regulation is missing from the other two RAF isoforms. This indicates that, as the most active RAF isoform, BRAF has acquired an additional regulatory layer absent from ARAF and CRAF. Within this framework, NRAS binds to the BRAF N-terminal regulatory assembly with sub-micromolar affinity (*K*_D_ ∼560 nM), indicating an isoform hierarchy of BRAF engagement (KRAS > NRAS >> HRAS) and reflecting a differential ability to overcome BSR-mediated gating^19^. The BSR-CRD interface thus emerges as a tunable regulatory brake on RAS-BRAF complex formation and as a potential vulnerability.

Our data support that oncogenic RAS mutations significantly strengthen interaction with the intact BRAF N-terminal assembly. Mutations at codons G13 and Q61 produced the largest increases in affinity, with several KRAS and NRAS mutants exhibiting low nanomolar binding to BRAF NT1. Notably, these enhanced interactions approached the affinity range typically associated with isolated RBD-mediated engagement, suggesting that oncogenic mutations partially overcome the regulatory constraints imposed by the BRAF N-terminal module. The cellular NanoBiT assays further supported the biophysical measurements. These findings provide a mechanistic explanation for enhanced RAF recruitment and sustained MAPK pathway activation observed in oncogenic RAS-driven cancers.

In addition to the differences observed in equilibrium affinity between KRAS and NRAS, further distinctions were evident at the level of kinetic behavior. KRAS consistently displayed slower dissociation kinetics relative to NRAS, indicating formation of more kinetically stable complexes than NRAS. Whereas oncogenic NRAS variants primarily exhibited enhanced association rates with relatively preserved dissociation kinetics, oncogenic KRAS mutants displayed both accelerated association and markedly reduced dissociation rates, producing substantially more stable KRAS-BRAF complexes. These findings suggest that KRAS and NRAS engage the BRAF regulatory region through distinct kinetic regimes, with KRAS favoring prolonged effector engagement and NRAS exhibiting more dynamic interaction behavior.

Our inhibitor profiling studies revealed substantial isoform- and mutation-specific differences in disruption of RAS-BRAF interactions. Among the compounds tested, the RAS(ON) inhibitor RMC6236 produced the most consistent suppression across both KRAS- and NRAS-driven complexes, including Q61 mutant variants. In contrast, switch-II pocket inhibitors displayed variable efficacy depending on both RAS isoform and mutational status. Adagrasib effectively disrupted KRAS-BRAF binding, producing an approximately 120-fold reduction in affinity by BLI and substantial inhibition in NanoBiT assays. However, its effects on NRAS-BRAF interactions were considerably weaker, with only modest reductions in affinity and complex stability. These findings are consistent with structural studies demonstrating that adagrasib binding is optimized for the KRAS switch-II pocket through interactions involving residues such as His95^36^, which are absent in NRAS. In contrast, Sotorasib displayed weaker effects, consistent with differences in switch-II pocket engagement shaped by H95/Y96 in KRAS^36^.

Mutations affecting codon 61 conferred particularly strong resistance to switch-II pocket inhibitors. KRAS(Q61R) and NRAS(Q61L/K) maintained substantial BRAF engagement despite inhibitor treatment, consistent with stabilization of the active GTP-bound conformation and reduced accessibility of the switch-II pocket. Similarly, BI-2865 showed only partial disruption of wild-type RAS-BRAF interactions and limited efficacy against Q61 mutants, likely reflecting its preferential binding to inactive GDP-bound conformations. These findings collectively suggest that the conformational state of RAS is a major determinant of inhibitor sensitivity and that mutations favoring constitutive activation can substantially impair switch-II pocket targeting.

Importantly, our BLI analyses further demonstrated that inhibitor-induced disruption of RAS-BRAF interactions occurs through distinct kinetic mechanisms. Adagrasib strongly destabilized KRAS–BRAF complexes by markedly accelerating dissociation kinetics, whereas its effects on NRAS complexes were comparatively limited. In contrast, BI-2865 produced a more moderate but broadly distributed reduction in binding across both isoforms. These observations indicate that disruption of RAS effector engagement depends not only on inhibitor occupancy but also on the extent to which inhibitor binding reshapes the conformational ensemble governing RAS-RAF complex stability.

Overall, our findings support a model in which the BRAF N-terminal regulatory region functions as a conformational gate that modulates RAS engagement, while intrinsic isoform-specific kinetic properties and oncogenic mutations determine the efficiency and stability with which this gate is overcome. Within this framework, oncogenic RAS mutations enhance signaling output by promoting more efficient and prolonged BRAF recruitment, whereas pharmacological inhibitors differentially disrupt these interactions depending on isoform-specific structural features and conformational state preferences. These results provide mechanistic insight into the dynamic regulation of RAS-RAF interactions and highlight how mutation-dependent changes in binding kinetics may contribute to differential therapeutic responses in RAS-driven cancers.

## Methods and Materials

### Plasmids

GST-KRAS/NRAS was created with standard Gibson Assembly (NEB) procedures with pGEX2T as the vector and the following primers: 5’ATCTGGTTCCGCGTGGATCCACTGAATATAAACTTGTGGTAG 3’ (GST-KRAS_For), 5’CAGTCAGTCACGATGAATTCTTACATAATTACACACTTTGTCTTTG 3’ (GST-KRAS_Rev), 5’ ATCTGGTTCCGCGTGGATCCACTGAGTACAAACTGGTG 3’ (GST-NRAS_For), 5’CAGTCAGTCACGATGAATTCTTACATCAACCACACATGG 3’(GST-NRAS_Rev), 5’GAATTCATCGTGACTGACTGACG 3’ (GST-vector_For), 5’GGATCCACGCGGAACCAG 3’ (GST-vector_Rev), and the variants of KRAS G12D,KRAS G12V,KRAS G13D,KRAS Q61L,KRAS Q61R and NRAS G12D, NRAS Q61L, NRAS Q61R, NRAS Q61K was generated by mutagenesis. His/MBP-BRAF NTs were created with standard Gibson Assembly (NEB) procedure in the pET28-MBP vector from the following primers: 5’ CATATGCTCGGATCCGCGGCGCTGAGCGGTG3’ (BRAF-RBD_For 1), 5’ CATATGCTCGGATCCTCACCACAAAAACCTATCGTTAG3’ (BRAF-RBD_For 151), 5’ GTTGTAAGAATTCAAGCTTACAACACTTCCACATGCAATTC3’ (BRAF-RBD_Rev 227), 5’ CAAAGAACTGAATTCAAGCTTACAAATCAAGTTGGT3’ (BRAF-RBD_Rev 288).

### Protein expression

Plasmids for all protein constructs were transformed into BL21 codon+ *E. coli* and cultured in LB broth until an OD600 0.6-0.8 was reached. For the BRAF NT constructs, the growth medium was supplemented with 100 μM ZnCl2. Following this, protein expression was induced by the addition of 0.4 mM IPTG, and the cultures were maintained overnight at 18 °C with constant shaking at 210 rpm. Finally, the cells were harvested by centrifugation, flash-frozen, and stored at −80 °C

### Protein purification

The pellets containing GST-tagged full-length KRAS/NRAS proteins were thawed and resuspended in lysis buffer (20 mM HEPES pH 7.4, 150 mM NaCl, 1 mM EDTA, 5% glycerol, and protease inhibitors). The whole-cell lysate was passed through the French pressure cell press at 1450 psi, followed by centrifugation to isolate the soluble fraction. This lysate was then incubated with pre-equilibrated glutathione resin for one hour at 4 °C. After extensive washing, the proteins were eluted using an elution buffer supplemented with 10 mM reduced glutathione. To facilitate nucleotide exchange, the eluted RAS was incubated in HEPES buffer with 10 mM EDTA and a 10-fold molar excess of GMPPNP for 30 minutes at 4 °C to promote the dissociation of bound nucleotides. Subsequent stabilization and rebinding were achieved by adjusting the MgCl2 concentration to 1 mM and rotating the mixture for 2 hours at 4 °C.

For BRAF-NT purification, cell pellets were resuspended in lysis buffer containing 20 mM HEPES (pH 8.0), 150 mM NaCl, 5% glycerol, and protease inhibitors. The soluble lysate was incubated with pre-equilibrated Ni-NTA resin for 1 h at 4 °C, washed extensively, and the His-tagged BRAF-NT protein was eluted using buffer containing increasing concentrations of imidazole (90-400 mM) in 20 mM HEPES (pH 7.4), 150 mM NaCl, and 5% glycerol. Both RAS and BRAF-NT proteins were further purified using size-exclusion chromatography on a Superdex 200 10/300 GL column. Purification of each stage was monitored via SDS-PAGE and Coomassie staining. Finally, the purified proteins were concentrated, flash-frozen in aliquots, and stored at -80 °C.

### Pulldowns

Proteins were combined in an 1:1 molar ratio and incubated in a binding buffer containing 20 mM HEPES (pH 7.4), 150 mM NaCl, 5% glycerol, and 0.125 mg/mL BSA. This mixture was subsequently incubated with 20 μL of glutathione Sepharose 4B resin (Cytiva) to facilitate the capture of GST-tagged complexes. Following incubation, the resins were subjected to stringent washing with a high-salt wash buffer (20 mM HEPES [pH 7.4], 500 mM NaCl, 5% glycerol, and 0.125 mg/mL BSA) to eliminate non-specifically bound proteins. Bound complexes were eluted by the addition of 30 μL of 4× SDS loading dye directly to the resin followed by thermal denaturation. Samples were subsequently resolved by SDS-PAGE, transferred onto nitrocellulose membranes, and analyzed by immunoblotting using a LI-COR imaging system.

### Surface plasma resonance (OpenSPR)

Interaction between RAS-BRAF: BRAF-NT1-4 constructs were immobilized on a Ni-NTA sensor chip to a minimum response of ∼2500 RU using immobilization buffer (20 mM HEPES pH 7.4, 150 mM NaCl, 0.05% Tween-20) according to the manufacturer’s protocol. Following immobilization, experiments were performed in running buffer containing 20 mM HEPES pH 7.4, 150 mM NaCl, 0.05% Tween-20, and 1% BSA, with glycerol levels matched to the RAS protein preparations. NRAS or KRAS analytes were flowed over the sensor surface at increasing concentrations ranging from 12.3 to 3000 nM at a flow rate of 30 μL/min for 10 min. Sensor surfaces were regenerated between cycles using 10 mM NaOH at 150 μL/min, followed by 3-4 min baseline stabilization. Binding kinetics were obtained by fitting the sensorgrams to a 1:1 interaction model using TraceDrawer, and reported values represent the mean ± standard deviation.

### Biolayer Interferometry (BLI)

Binding interactions between BRAF NT1 and RAS proteins were measured using biolayer interferometry on an Octet N1. His-tagged BRAF NT1 was immobilized on Anti-Penta-HIS (HIS1K) biosensors by loading 4 μL of protein (10-35 ng/μL) to achieve an immobilization response of ∼1.0-1.2 nm. RAS proteins were pre-incubated with 10 µM compounds for 2 h prior to analysis. Experiments were conducted in buffer containing 20 mM HEPES (pH 7.4), 150 mM NaCl, 0.05% Tween-20, with glycerol levels matched to the RAS protein preparations. After loading (3-4 min, 2000 rpm) and baseline equilibration, KRAS and NRAS mutants were introduced over a concentration range of 37 nM to 12 μM to monitor association, followed by dissociation in assay buffer. Binding data were globally fitted using a 1:1 binding model in Octet N1.4 analysis software to obtain kinetic parameters and equilibrium dissociation constants (*K_D_*), with curves plotted in GraphPad Prism. Measurements represent at least three independent replicates, and reported *K_D_* values correspond to the mean ± standard deviation.

### NanoBiT Constructs Cloning

NanoBiT® CMV MCS BiBiT vector, which contains a BRAF N- terminal fused with LgBiT and CRAF N-terminal fused with SmBiT, were purchased from Promega. KRAS^LgBiT^ and BRAF^SmBiT^ was generated using the standard Gibson Assembly (NEB HiFi Assembly) using NanoBiT® BiBiT as the vector and following primers: 5’ CTCCGCCCCCCAGCGACACCATAATTACACACTTTGTCTTTG 3’, 5’ CCACAAGTTTATATTCAGTCATGGCGATCGCTAGCTGCAAA 3’, 5’ GGCGATCGCTAGCTGCAAAAAGAACAAG 3’, 5’ TCCGCCCCCCAGCGACAC 3’, and NRAS^LgBiT^ and BRAF^SmBiT^ was generated using the standard Gibson Assembly using NanoBiT® BiBiT as the vector and following primers: 5’ CCTCCGCCCCCCAGCGACACCATCACCACACATGGCAATC 3’, 5’ CACCAGTTTGTACTCAGTCATGGCGATCGCTAGCTGCAAAA 3’, 5’ GGCGATCGCTAGCTGCAAAAAGAACAAG 3’.

### NanoBiT Luminescence Assay

HEK293 cells were transiently transfected with 2 µg per well of wild-type full-length BRAF^SmBiT^-KRAS^LgBiT^, WTFL BRAF^SmBiT^-NRAS^LgBiT^ or KRAS/NRAS mutants and incubated at 37 °C. Twenty-four hours later, cells were trypsinized, pelleted, and resuspended in OptiMEM containing 4% FBS, then re-seeded onto 96-well plates at 6×10⁴ cells per well and incubated overnight. Cells were treated with 10 µM of Sotorasib, Adagrasib, BI 2865, or RMC 6236 for 4 hours. Following treatment, 10 µM furimazine substrate was added, and luminescence was measured at 470-480 nm using a CLARIOstar plate reader. Data were analyzed for statistical significance using paired t-tests where appropriate, followed by Tukey’s post-hoc test. P-values are indicated as *P<0.05, **P<0.01, ***P<0.001. All graphs show mean ± SD with individual biological replicate data points.

### Antibodies used in immunoblots

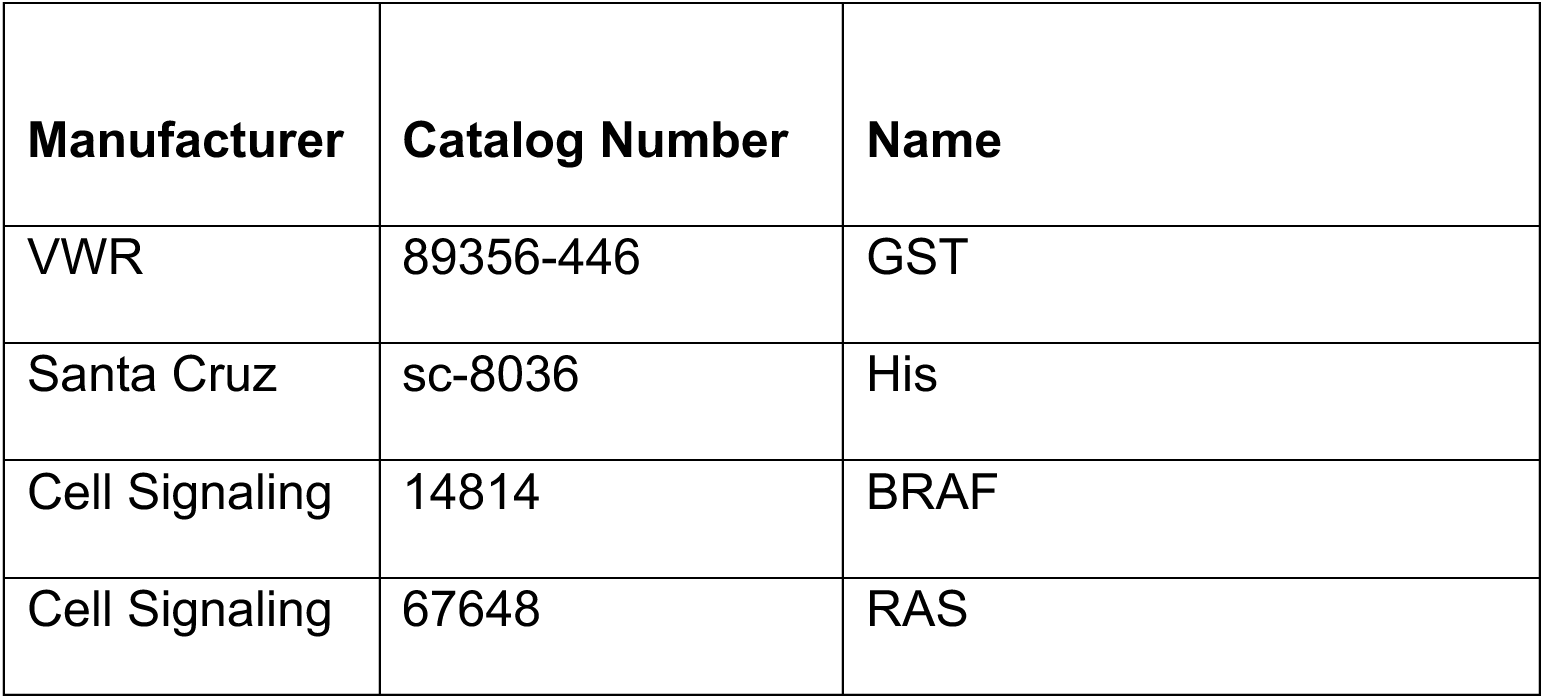
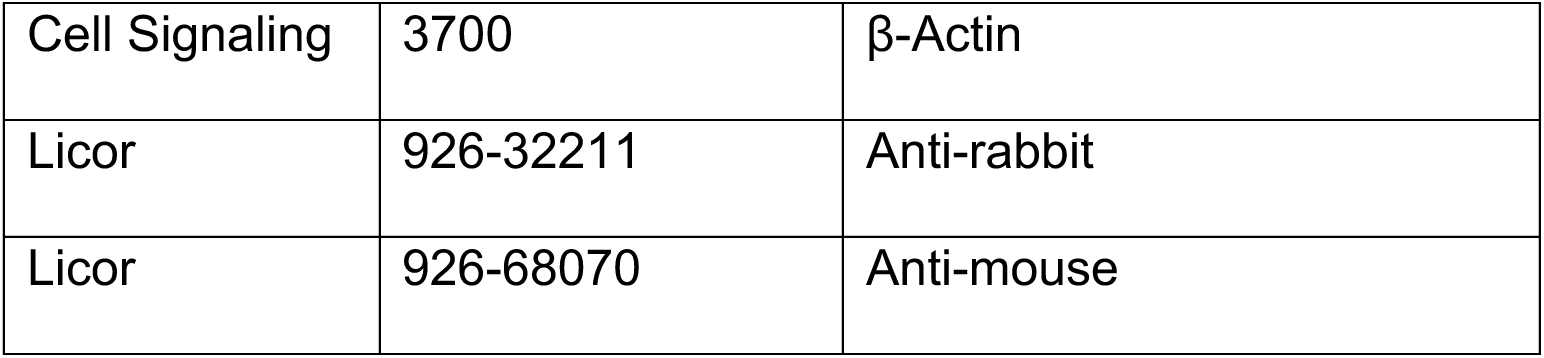

### Inhibitors

BI-2865 (E1474), RMC-6236 (E1597), Sotorasib (S8830), and Adagrasib (S8884) were purchased from Selleckchem.

### Funding Sources

The authors declare their funding sources as follows: NIH funding R01GM138671 to ZW.

### Data Sharing Plan

All data are included in the manuscript and supporting information.

## Supplementary Information

**Figure S1.**
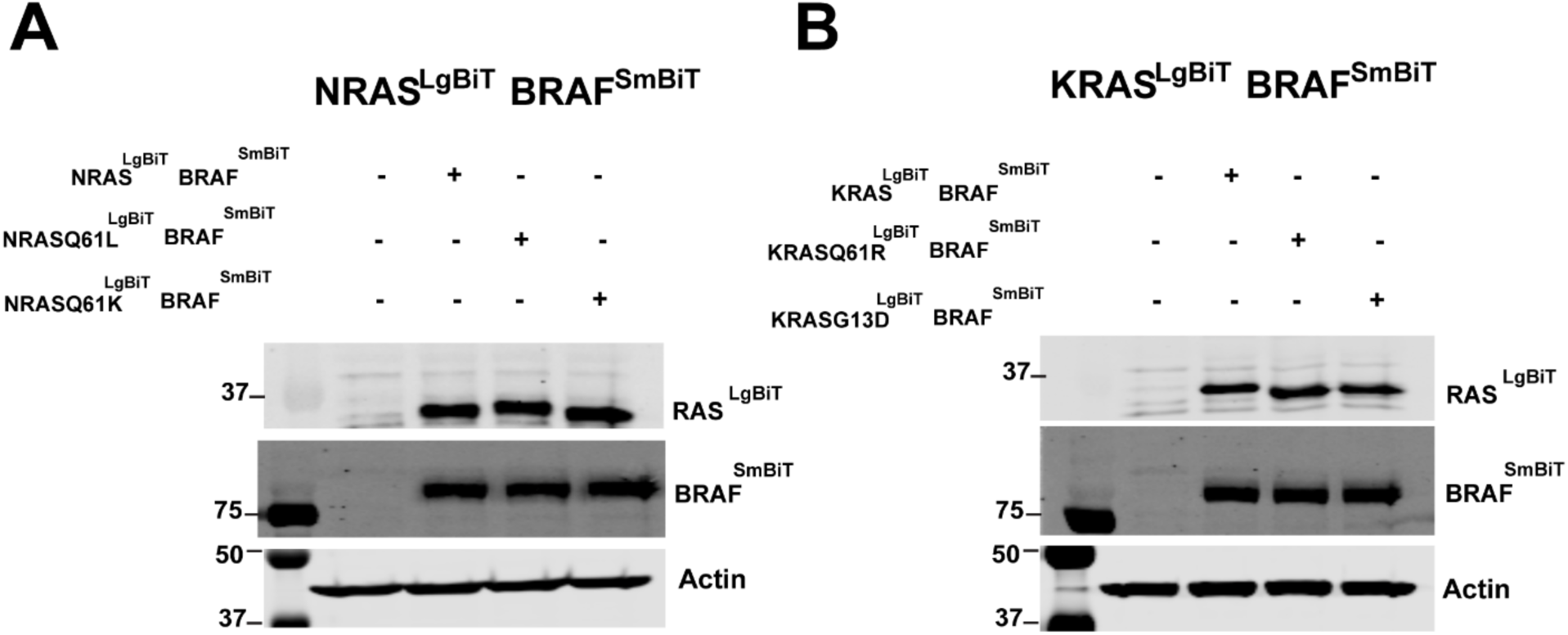
RAS-BRAF protein expression in cells. A) Representative immunoblots of HEK293 cells expressing NRAS^LgBiT^ (WT, Q61L, Q61K) with BRAF^SmBiT^. (B) Representative immunoblots of HEK293 cells expressing KRAS^LgBiT^ (WT, Q61R, G13D) with BRAF^SmBiT^. HEK293 Cells were transfected and lysed 48h post-transfection. Protein expression was confirmed by Western blotting. β-actin was used as a loading control. Experiments were performed in three independent biological replicates.

**Figure S2.**
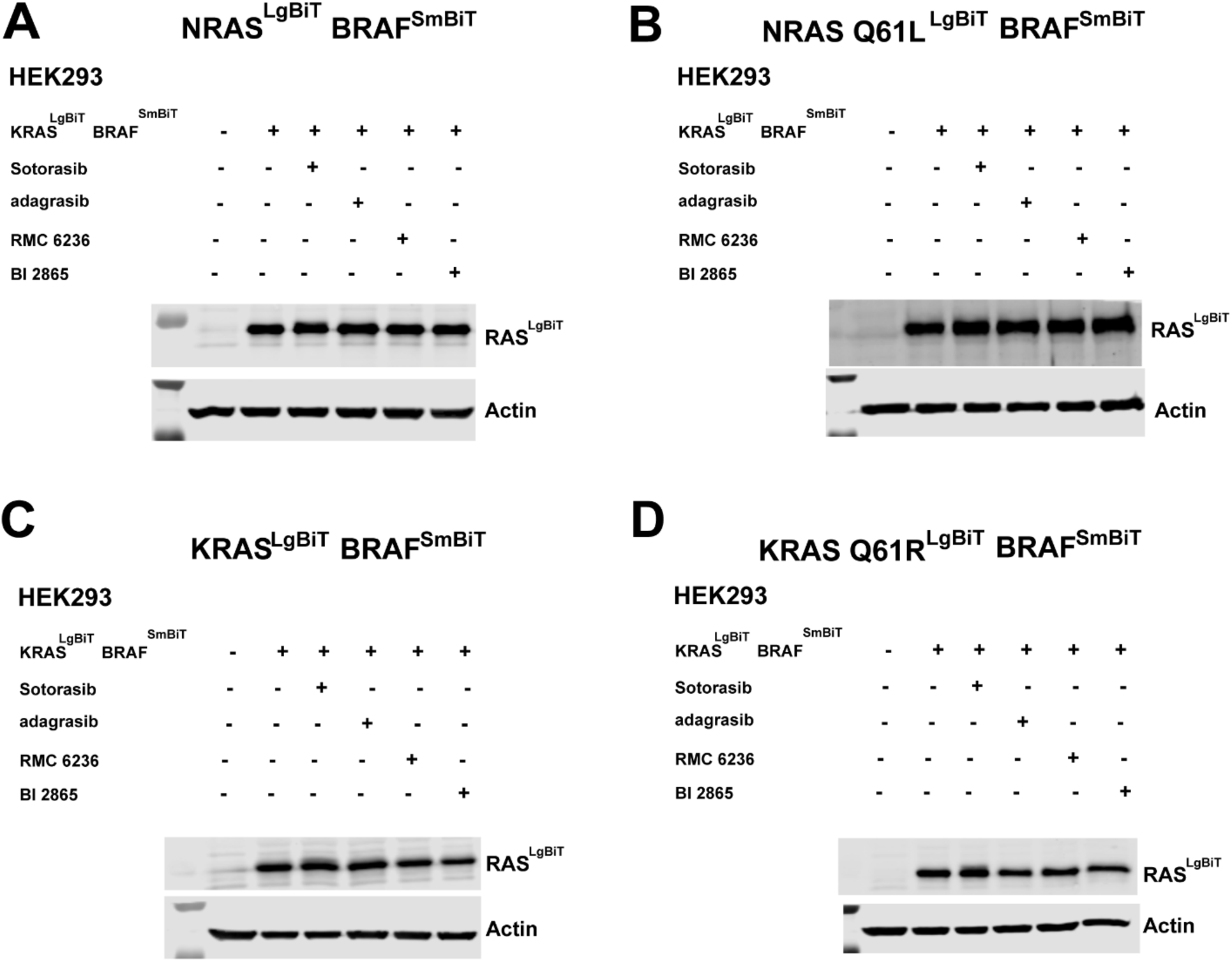
RAS inhibitors disrupt RAS-BRAF interactions without altering protein expression in cells. A-B) Representative immunoblots of HEK293 cells expressing (A) NRAS^LgBiT^ with BRAF^SmBiT^ (B) NRASQ61L^LgBiT^ with BRAF^SmBiT^ in the absence or presence of the indicated RAS inhibitors (Sotorasib, Adagrasib, RMC 6236, and BI 2865). C-D) Immunoblots of HEK293 cells expressing (C) KRAS^LgBiT^ with BRAF^SmBiT^ or (D) KRASQ61R^LgBiT^ with BRAF^SmBiT^ under the same treatment conditions. All compounds were used at 10 µM with a 4 h incubation prior to BLI measurements. Experiments were performed in two independent biological replicates.

**Figure S3.**
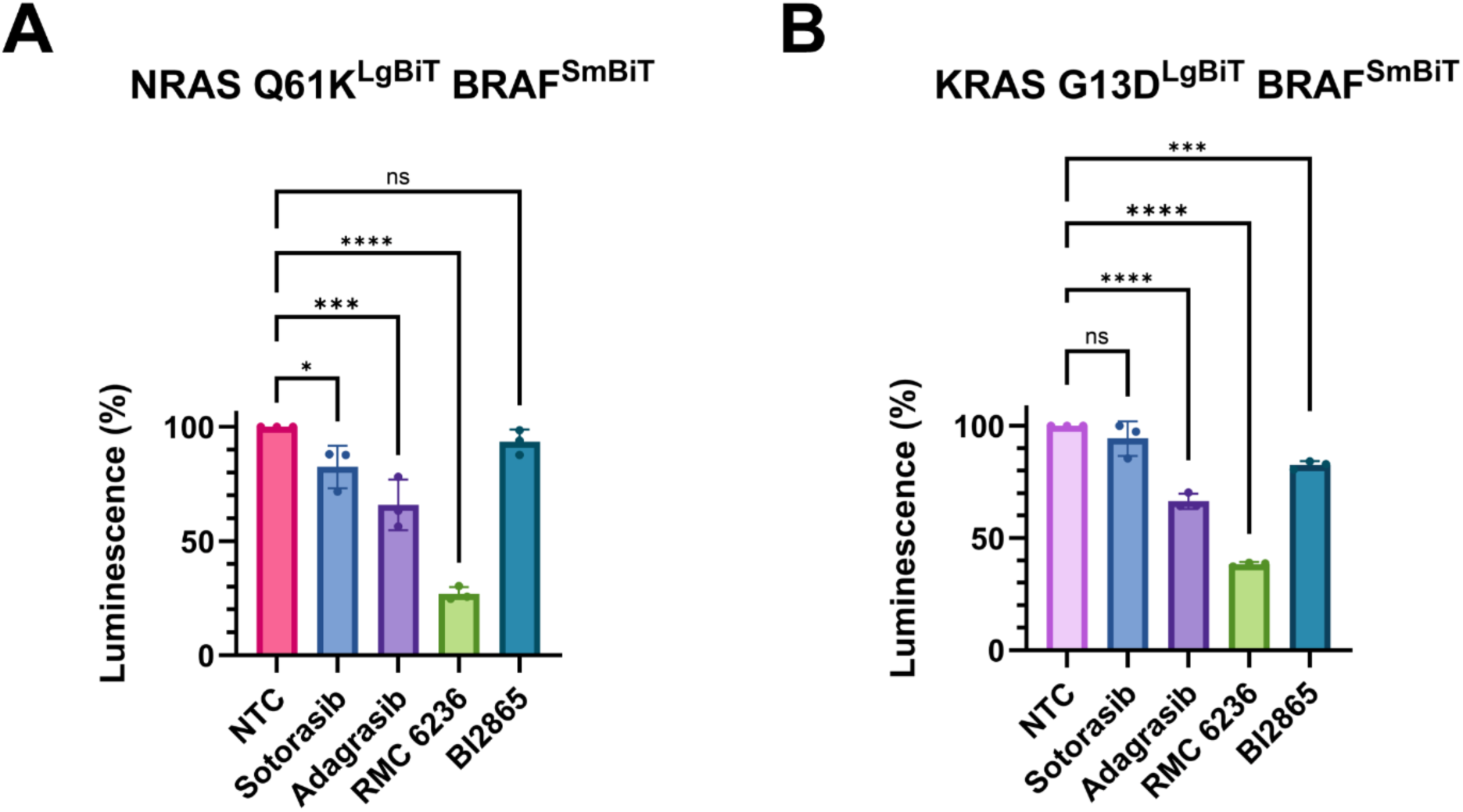
Mutation-specific regulation of RAS-BRAF interactions and inhibitor sensitivity in cells. (A-B) Effects of RAS inhibitors (sotorasib, adagrasib, RMC6236, and BI2865) on (A) NRAS Q61K^LgBiT^ with BRAF^SmBiT^(B) KRAS G13D^LgBiT^ with BRAF^SmBiT^ interactions measured by NanoBiT luminescence. HEK293 cells were treated with 10 µM of each compound for 4h prior to analysis. Data were obtained from three independent biological replicates, each performed in technical triplicate.

**Table S1.** OpenSPR kinetic parameters for NRAS binding to BRAF NT1–NT4.

| Ligand | Analyte | $k_{on}$<br>(1/Ms) | $k_{off}$ (1/s) | $K_D$<br>(nM) | Avg<br>$K_D$<br>(nM) | St<br>dev<br>(nM) | BI<br>(signal<br>RU) | Bmax | Chi2<br>value | U-value:<br>kd (%) |
| --- | --- | --- | --- | --- | --- | --- | --- | --- | --- | --- |
| NT1 1-288<br>(BSR+RBD+CRD) | NRAS | 6050 | 0.00378 | 624 | 587 | 52.3 | 0.10 | 146.4 | 14.05 | 3.80 |
| NT1 1-288<br>(BSR+RBD+CRD) | NRAS | 10100 | 0.00553 | 550 |  |  | 0.10 | 106.5 | 5.14 | 1.90 |
| NT2 1-227<br>(BSR+RBD) | NRAS | 34800 | 0.00328 | 94 | 85 | 12.7 | 0.10 | 158.0 | 10.10 | 1.20 |
| NT2 1-227<br>(BSR+RBD) | NRAS | 30000 | 0.00229 | 76 |  |  | 0.28 | 734.0 | 25.51 | 1.70 |
| NT3 151-288<br>(RBD+CRD) | NRAS | 37600 | 0.00201 | 54 | 60 | 9.8 | 0.10 | 190.6 | 19.30 | 1.70 |
| NT3 151-288<br>(RBD+CRD) | NRAS | 45700 | 0.00306 | 67 |  |  | 0.10 | 103.3 | 10.61 | 2.10 |
| NT4 151-227<br>(RBD) | NRAS | 28300 | 0.00304 | 108 | 94 |  | 0.10 | 347.4 | 33.58 | 1.90 |
| NT4 151-227<br>(RBD) | NRAS | 57200 | 0.00452 | 79 |  |  | 0.10 | 462.1 | 22.86 | 2.20 |

**Table S2.** OpenSPR kinetic parameters for NRAS mutants binding to BRAF NT1.

| Ligand | Analyte | $k_{on}$<br>(1/Ms) | $k_{off}$<br>(1/s) | $K_D$<br>(nM) | Avg<br>$K_D$<br>(nM) | St<br>dev<br>(nM) | BI<br>(signal<br>RU) | Bmax | Chi2<br>value | U-value:<br>kd (%) |
| --- | --- | --- | --- | --- | --- | --- | --- | --- | --- | --- |
| NT1 1-288<br>(BSR+RBD+CRD) | NRAS<br>Q61L | 37500 | 0.00341 | 91 | 87 | 5.7 | 0.10 | 233.6 | 18.11 | 1.70 |
| NT1 1-288<br>(BSR+RBD+CRD) | NRAS<br>Q61L | 37200 | 0.00310 | 83 |  |  | 0.10 | 206.2 | 10.99 | 1.60 |
| NT1 1-288<br>(BSR+RBD+CRD) | NRAS<br>Q61R | 44300 | 0.00376 | 85 | 91 | 8.1 | 0.10 | 170.1 | 8.83 | 1.40 |
| NT1 1-288<br>(BSR+RBD+CRD) | NRAS<br>Q61R | 41700 | 0.00403 | 97 |  |  | 0.10 | 153.1 | 18.10 | 2.20 |
| NT1 1-288<br>(BSR+RBD+CRD) | NRAS<br>Q61K | 28500 | 0.00388 | 136 | 116 | 28.3 | 0.10 | 121.1 | 6.03 | 2.10 |
| NT1 1-288<br>(BSR+RBD+CRD) | NRAS<br>Q61K | 38500 | 0.00370 | 96 |  |  | 0.10 | 160.1 | 30.00 | 2.71 |
| NT1 1-288<br>(BSR+RBD+CRD) | NRAS<br>G12D | 33700 | 0.00384 | 114 | 116 | 2.5 | 0.10 | 118.3 | 14.93 | 2.20 |
| NT1 1-288<br>(BSR+RBD+CRD) | NRAS<br>G12D | 32400 | 0.00385 | 119 |  |  | 0.10 | 267.9 | 16.99 | 1.70 |

**Table S3.** OpenSPR kinetic parameters for KRAS mutant binding to BRAF NT1.

| Ligand | Analyte | $K_{on}$<br>(1/Ms) | $K_{off}$<br>(1/s) | $K_D$<br>(nM) | Avg<br>$K_D$<br>(nM) | St<br>dev<br>(nM) | BI<br>(signal<br>RU) | Bmax | Chi2<br>value | U-value:<br>kd (%) |
| --- | --- | --- | --- | --- | --- | --- | --- | --- | --- | --- |
| NT1 1-288<br>(BSR+RBD+CRD) | KRAS<br>Q61L | 30300 | 0.00063 | 21 | 20 | 1.3 | 0.10 | 233.5 | 18.11 | 1.70 |
| NT1 1-288<br>(BSR+RBD+CRD) | KRAS<br>Q61L | 41800 | 0.00078 | 19 |  |  | 0.10 | 206.2 | 10.99 | 1.60 |
| NT1 1-288<br>(BSR+RBD+CRD) | KRAS<br>Q61R | 6540 | 0.00032 | 49 | 38 | 15.7 | 0.10 | 170.1 | 8.83 | 1.40 |
| NT1 1-288<br>(BSR+RBD+CRD) | KRAS<br>Q61R | 8370 | 0.00022 | 27 |  |  | 0.10 | 153.1 | 18.10 | 2.20 |
| NT1 1-288<br>(BSR+RBD+CRD) | KRAS<br>G13D | 25700 | 0.00062 | 23 | 25 | 1.9 | 0.10 | 121.1 | 6.03 | 2.10 |
| NT1 1-288<br>(BSR+RBD+CRD) | KRAS<br>G13D | 30500 | 0.00081 | 26 |  |  | 0.10 | 160.0 | 30.00 | 2.70 |
| NT1 1-288<br>(BSR+RBD+CRD) | KRAS<br>G12D | 8120 | 0.00174 | 214 | 204 | 14.8 | 0.10 | 118.1 | 14.93 | 2.20 |
| NT1 1-288<br>(BSR+RBD+CRD) | KRAS<br>G12D | 9850 | 0.00190 | 193 |  |  | 0.10 | 267.9 | 16.99 | 1.70 |

**Table S4.** BLI Kinetic Analysis of RAS-BRAF NT1 Interactions with inhibitors.

| **Binding Partners** | **K<sub>D</sub> (μM)** | **K<sub>on</sub> (1/Ms)** | **K<sub>off</sub> (1/s)** | **χ<sup>2</sup>** | **R<sup>2</sup>** |
| --- | --- | --- | --- | --- | --- |
| BRAF NT1-KRAS+Adagrasib | 21.91 | 1.022 x 10<sup>4</sup> | 2.239 x 10<sup>-1</sup> | 1.03 | 0.988 |
|  | 28.01 | 9.069x 10<sup>3</sup> | 2.546 x 10<sup>-1</sup> | 2.21 | 0.976 |
| BRAF NT1-KRAS+BI2865 | 1.33 | 3.425x 10<sup>3</sup> | 4.561 x 10<sup>-3</sup> | 0.32 | 0.992 |
|  | 1.52 | 4.708 x 10<sup>3</sup> | 7.154 x 10<sup>-3</sup> | 0.62 | 0.984 |
| BRAF NT1-NRAS+Adagrasib | 1.83 | 3.184 x 10<sup>4</sup> | 5.832 x 10<sup>-2</sup> | 1.01 | 0.984 |
|  | 1.75 | 3.521 x 10<sup>4</sup> | 6.149 x 10<sup>-2</sup> | 0.56 | 0.990 |
| BRAF NT1 -<br>NRAS+ BI2865 | 1.19 | 3.564 x 10<sup>3</sup> | 4.227 x 10<sup>-3</sup> | 0.36 | 0.993 |
|  | 1.23 | 2.984 x 10<sup>3</sup> | 3.669 x 10<sup>-3</sup> | 0.46 | 0.985 |

